# Leaf side determines the relative importance of dispersal versus host filtering in the phyllosphere microbiome

**DOI:** 10.1101/2022.08.16.504148

**Authors:** Wenke Smets, Mason K Chock, Corinne M Walsh, Caihong Qiu Vanderburgh, Ethan Kau, Steve E Lindow, Noah Fierer, Britt Koskella

## Abstract

Leaf surface-associated bacterial communities play a significant role in plant health and have therefore been the focus of increasing interest. Despite this, we currently lack a predictive understanding of how leaf-associated bacterial communities are structured within and across hosts, including how leaf traits shape this variation and how community assembly processes may differ across distinct microbial habitats on a leaf. In this study, we characterize the composition of bacterial phyllosphere communities from the upper and lower leaf surfaces of 66 plants across 24 species grown at a common site using 16S rRNA amplicon sequencing. By comparing leaves that vary in pH and stomatal densities and analyzing the leaf surfaces separately, we were able to test the key factors shaping the phyllosphere across host plant species from diverse geographical origins. We found a surprisingly large shared/core microbiome across species, as well as a strong effect of plant species and native origin in shaping composition. Importantly, we found the lower leaf side, where pH values are generally lower and stomatal densities higher, to have lower taxonomic richness relative to the upper leaf side. While the upper leaf side community appears to be more strongly influenced by dispersal effects, the lower leaf appears to be more strongly influenced by plant host filtering effects, as supported by higher relative abundance of shared core taxa and higher signatures of endemism. This work highlights important differences in community assembly processes across the upper and lower leaf microbiomes and underscores the importance of considering differences among habitats within a host when explaining microbial community assembly and composition.

## Introduction

Phyllosphere (leaf-associated) bacterial communities are increasingly recognized for their role in plant health, modulating chemical emissions of plants, providing nutrients to plants (e.g. nitrogen fixation) and protecting against plant pathogens (1–3). Leaves typically harbor bacterial populations ranging in size from 10^6^ to 10^7^ cells/cm^2^ leaf (4). Since aerial plant compartments are estimated to comprise 60% of the global plant biomass, the phyllosphere microbiome contributes substantially to global biogeochemical cycles and ecosystems worldwide (5). The bacterial communities found on leaves are distinct from those in other environmental habitats or host organisms, but also differ substantially in composition among host plant species. Some of the distinctiveness of epiphytic bacterial communities can be attributed to abiotic factors, including UV exposure (6, 7), temperature (8), and the differential availability of liquid water (9, 10). Likewise, the composition of leaf-associated microbial communities is driven by unique biotic factors, including leaf structure and physiology (11, 12), differential exposure to immigrant inoculum sources (13–16), and interactions among microbes, including with bacteriophage (17, 18). In addition to these deterministic processes, phyllosphere communities – like all microbiomes - are subject to stochasticity during assembly due to the random nature of arrival time and the order of arrival of dispersing microbes (15, 19).

Although phyllosphere community variation has been attributed to differences in plant species identity and phylogeny (20–22), even leaves from the same individual plant or those from different individual plants of a given plant species at a given sampling site can harbor remarkably distinct bacterial communities (23, 24). The relative importance of deterministic (host filtering) and neutral (dispersal) processes in shaping the microbiome remains controversial and has recently been shown to differ as a function of plant neighborhood composition and biomass (25), which might help explain the variation among published studies regarding the relative importance of plant selection on the phyllosphere microbiome (26, 27, 28). Moreover, the apparent importance of deterministic versus neutral processes could be related to the fact that it is often implicitly assumed that the leaf microbiome of an individual plant, or even a particular plant species, constitutes a single community.

Leaves of most plant species have distinct anatomies and physiologies between their upper (adaxial) and lower (abaxial) surfaces. Topographic features that differ between the leaf sides are expected to influence microbial community size and interactions, as exemplified by bacterial aggregations often observed in grooves between plant cells, or by trichomes and stomata (29). Topographical features also strongly influence the distribution and movement of water on leaves, thereby influencing bacterial localization and their association with water (10). Environmental stressors typically encountered in the phyllosphere, including UV irradiation, periodic desiccation, and variation of humidity (4, 7, 30), are expected to be more intense on the upper than on the lower leaf surface. Moreover, in most plant species, stomatal density is higher on the lower surface of the leaf than on the upper surface (31), possibly leading to higher local relative humidity due to the trapping of evapotranspiration within the laminar boundary layer surrounding the leaf (32). Such higher stomatal densities might lead to strong host ‘filtering’ effects on the lower leaf surfaces. In contrast, the upper surfaces of leaves are more likely to intercept airborne bacteria than the underside, due to higher rates of impaction and deposition of airborne particles (33–35). Such processes would decrease the relative strength of host filtering on microbiome composition on the upper leaf compared to the lower (16). Although observations comparing leaf surface bacterial communities are limited, measurements of viable cells and microscopic observations have found higher numbers of bacteria on the lower compared to upper surface of leaves in sun-exposed peanut plants and in *Arabidopsis* plants grown in the field (36, 37). We hypothesize that the higher humidity and elevated availability of resources make the lower surface of a leaf more conducive to bacterial growth, activity, and interaction with the host than the upper leaf surface. These factors, combined with reduced dispersal of microbes onto lower leaf surfaces, would lead to a stronger signature of host filtering for phyllosphere microbial communities on the lower surface versus the upper surface of leaves.

To test the relative impact of leaf physiology and differential exposure of leaves to bacterial immigration on the microbial community, we sampled replicate leaves from 24 plant species growing in a botanical garden in Berkeley, California. By comparing the composition of epiphytic bacterial communities between the upper and lower surface of leaves, we were able to directly test the hypotheses that (1) the influence of host plant species on leaf microbiome composition is higher on the underside of leaves and (2) upper leaf surfaces are more likely to harbor transient taxa that disperse and accumulate via atmospheric deposition, thereby overcoming host filtering effects on community composition.

## Results

We used 16S rRNA gene sequencing to characterize the phyllosphere communities found on 132 leaf samples representing 24 different plant species, with all plant species sampled in the University of California, Berkeley Botanical Garden. After quality filtering (see Methods), we obtained a total of 7976 Amplicon Sequence Variants (ASVs) representing 1068 bacterial genera. The most abundant genera were *Sphingomonas* (11.5%), *Methylobacterium* (7.5%), and *Hymenobacter* (6.3%), and the 11 most abundant families constituted between 43% and 83% of the total community of each plant species (Figure 1A). These families represented two thirds of all core ASVs, which were determined using the occupancy-abundance approach developed by Shade and Stopnisek (38). Here we define core ASVs as microbiome members shared across all 132 leaf samples (39). All 66 core ASVs identified in this study are listed in Supplemental Table 1. Despite having sampled the phyllosphere of 24 different plant species, this analysis uncovered a cross-species core microbiome that encompassed a considerable fraction of the communities, as core ASVs represented 42% of all reads.

**Figure 1:**
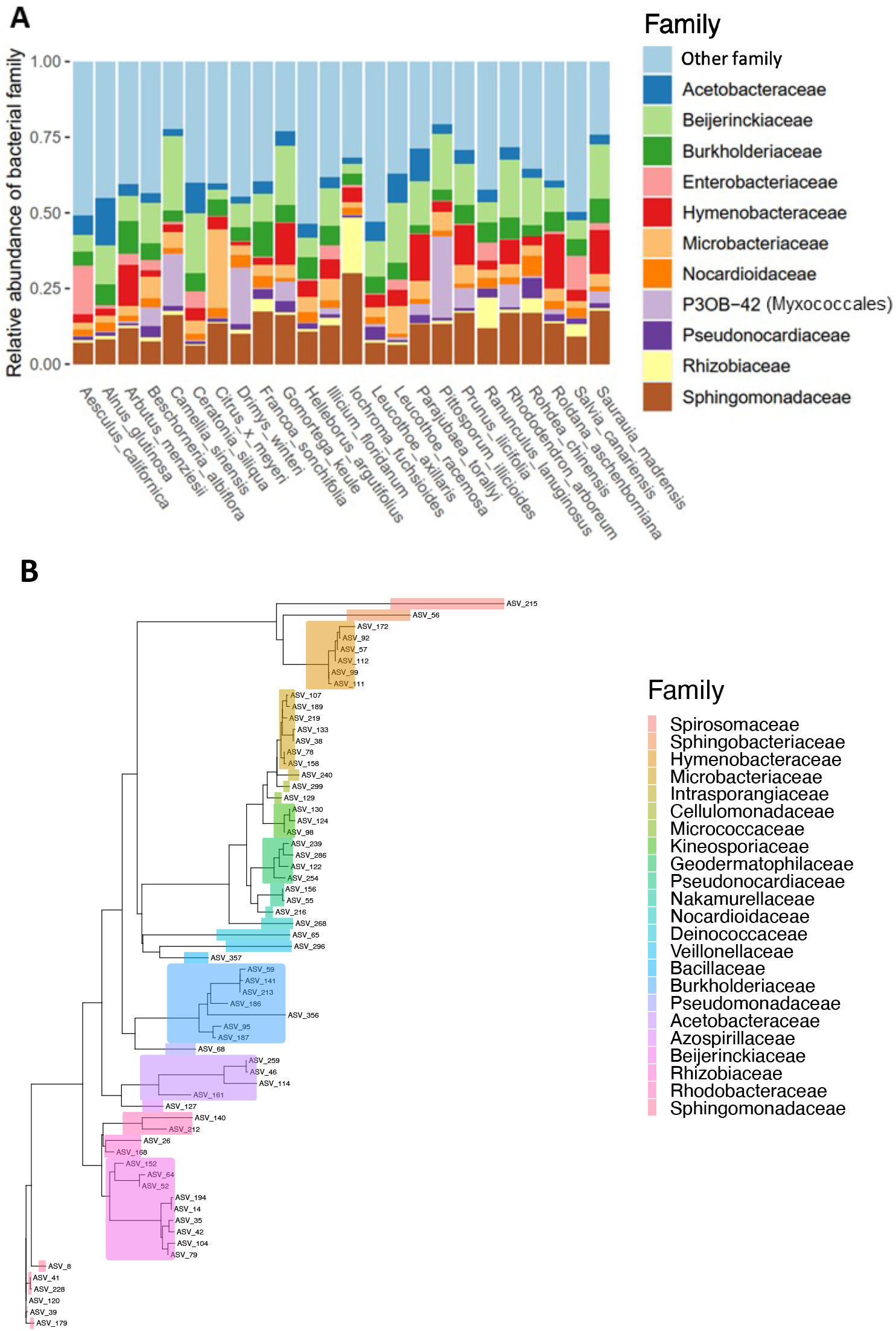
(A) Barplot representing the relative abundance of the 11 most abundant bacterial families. Each bar represents the average relative abundance in samples of a given plant species (generally n=6, representing upper and lower leaf surface samples from three individual plants). The residual taxa bar represents the relative abundance of the remaining less abundant families. (B) Phylogenetic tree of the 66 ‘core’ bacterial ASVs, determined using occupancy-abundance distributions. Key legend is in order of tips from top to bottom.

When compositions across phyllosphere communities were compared using Bray-Curtis dissimilarity analyses, an overall PERMANOVA model (including all samples) suggests that significant variation in microbiome composition is explained by, in order of signficance, Plant individual/leaf sample, Host plant species, Leaf side by Host plant species interaction, Leaf side, Stomatal density, and Leaf surface pH (Table 1). The strong interaction effect observed between leaf side and plant species (Table 1; R^2^ = 0.12; Supplemental Figure 1) motivated us to further analyze the upper and lower leaf samples separately given that, even within a given plant species, the two leaf sides appear to harbor distinct bacterial communities.

**Table 1:** R^2^ values for leaf and plant characteristics of interest explaining Bray-Curtis dissimilarity variation between samples in a PERMANOVA model (9999 permutations). Factors are in the order they were added to the model. The full model for all variables examined can be found in Supplemental Table 2.

| Factor | Overall | Upper surface | Lower surface |
| --- | --- | --- | --- |
| | $R^2$ | $R^2$ | $R^2$ |
| Leaf surface pH | 0.01** | 0.02 | 0.02 |
| Stomatal density | 0.02*** | 0.02* | 0.02* |
| Leaf side (upper/lower) | 0.02*** | NA | NA |
| Host plant species | 0.21*** | 0.28*** | 0.30*** |
| Plant individual | 0.32*** | NA | NA |
| Interaction of side and host plant species | 0.12*** | NA | NA |

$p < 0.05 = *$ , $p < 0.01 = **$ , $p < 0.001 = ***$

### Analyses of the upper and lower leaf surface microbiomes

Because leaf side explained such significant amount of variation in bacterial composition, we compared community characteristics between upper and lower leaf surfaces separately. For the upper leaf surface samples, host plant species (R^2^ = 0.28, p-value < 0.001) and region of native origin (R^2^ = 0.17, p-value < 0.001) remained the most important factors explaining community dissimilarities, while pH (R^2^ = 0.02, p-value = 0.08) and stomatal density (R^2^ = 0.02, p-value = 0.19) were no longer significant contributors (Table 1). The occupancy-abundance curve of the upper leaf surface samples’ fit to the neutral model using maximum likelihood estimation was 0.121 (Figure 2A), and showed a left hand skew (more taxa with lower abundance and higher occupancy) compared to that from lower leaf surface samples (Figure 2C). For the lower leaf surface samples, host plant species (R^2^ = 0.30, p-value < 0.001) and region of origin (R^2^ = 0.14, p-value < 0.001) were again the most important factors explaining community variation, with stomatal density (R^2^ = 0.02, p-value =0.03) as an additional significant contributor, but pH (R^2^ = 0.02, p-value = 0.09) again became non-significant. The fit of these observations to the neutral occupancy-abundance model was 0.045 for the lower leaf surface samples, suggesting stronger impacts of deterministic processes compared to the upper leaf (Figure 2A, B).

**Figure 2:**
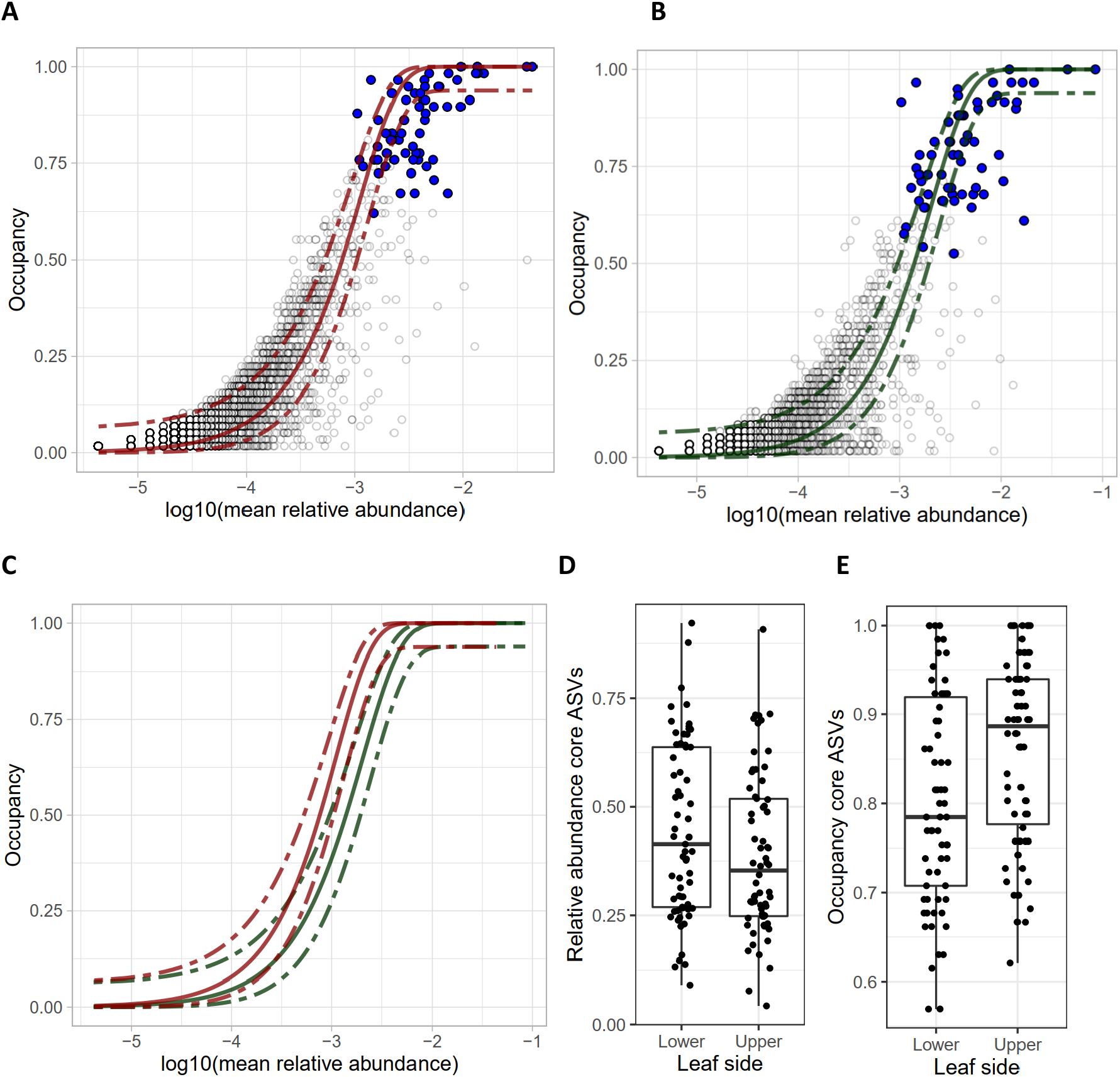
Occupancy-abundance curves of ASVs in (A) the upper leaf surface samples only, and (B) the lower leaf samples only. Each circle represents an ASV, with the blue dots representing core ASVs. The lines in each graph indicate the neutral model based on upper samples (green) and lower samples (blue), which are directly compared in (C). The (D) relative abundance and (E) occupancy of core ASVs for each of the leaf sides differed significantly (paired Wilcoxon p-value = 0.046 and p-value < 0.001, respectively).

### Alpha diversity differs significantly between leaf sides

Both species richness and Simpson’s index of the upper leaf surface phyllosphere communities were significantly higher than those of the lower leaf surface (Figure 3A, B). The weighted degree of endemism, an indication of how restricted ASVs of a sample are to one or several host plant species compared to the group, was significantly higher for the lower leaf surfaces than for the upper leaf surfaces (paired Wilcoxon p-value = 0.005; Figure 3C). This indicates a higher relative abundance of endemic ASVs on the lower surfaces of leaves than on the upper surfaces.

**Figure 3.**
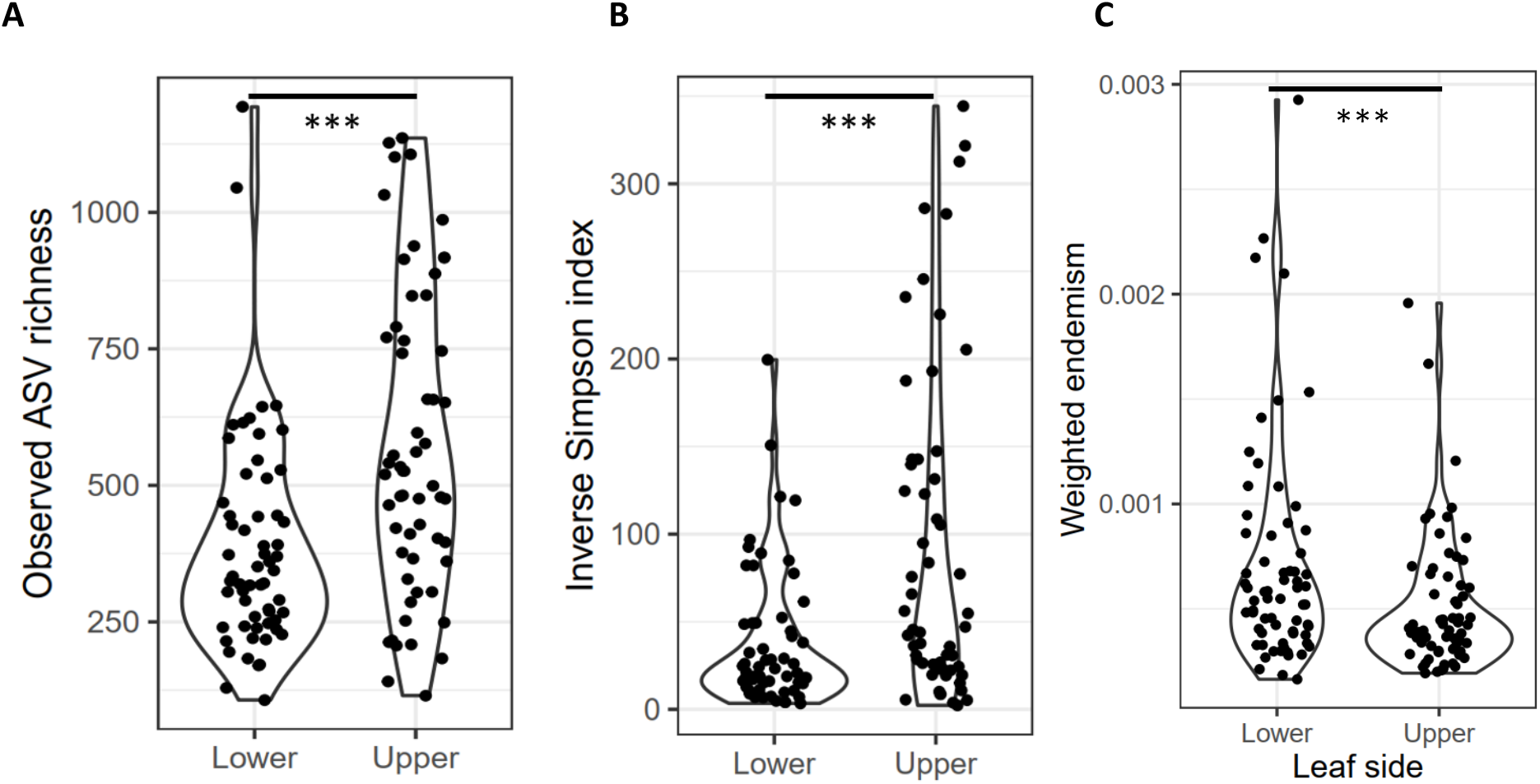
(A) The observed ASV richness and (B) the inverse simpson index were determined per sample and showed significantly lower alpha diversities for lower leaf surfaces than for upper leaf surfaces. (C) The weighted degree of endemism for samples of each leaf side.

A differential abundance analysis of ASVs by leaf side identified several taxa that were preferentially present on the upper or lower leaf surfaces (Figure 4B). We found that 67% of the ASVs significantly associated with the lower leaf surface belonged to the core phyllosphere microbiome found in this study (Supplemental table 1), whereas only 13% of the ASVs significantly associated with the upper leaf surface belonged to the core microbiome. This suggests that bacteria which are more likely to have a functional relationship with the plant host are also more likely to be found on the lower leaf surface. Moreover, core ASVs were found to occupy a lower proportion of ASVs overall, but had higher relative abundance on the lower leaf surface relative to the upper leaf surface (paired Wilcoxon p-value = 0.046 and p-value < 0.001, respectively; Figure 2D, E).

**Figure 4:**
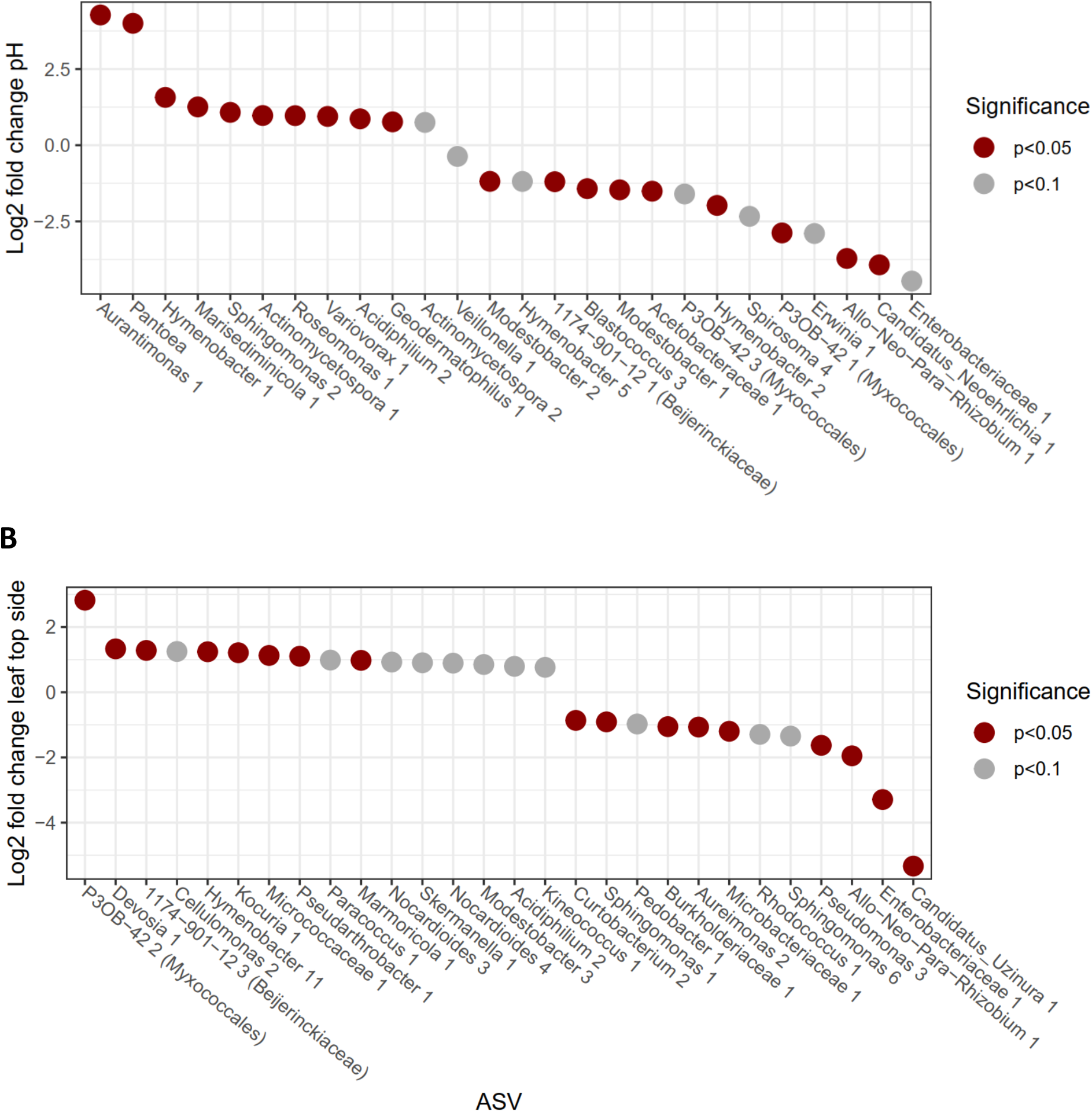
Significant and near-significant bacterial ASVs associated with various leaf traits. (A) Enrichment of taxa on more acidic leaf surface is represented by a postivie Log2 fold change while a negative log2 fold change indicate a higher abundance of bacterial taxa on on a more neutral leaf surface. (B) Differentially abundant taxa between upper and lower surface communities of leaves. A positive log2 fold change indicates a preference for the upper sides of the leaves, a negative log2 fold change indicates a preference for the lower sides of the leaves.

### Stomatal density of leaves impacts microbiome composition and diversity

Stomatal densities differed markedly between the upper and lower sides of the leaves of all species, as few or no stomata were found within the observed areas of the upper leaf surface of most plant species (Figure 5B). In the case of the lower leaf surface, where most stomata were found, stomatal density is negatively, but not significantly, correlated with pH (p-value = 0.14, tau = −0.23). We did, however, observe a significant negative correlation between stomatal density and ASV richness (p-value = 0.003, tau = −0.27) across the lower leaf surface samples. A differential abundance analysis considering only the lower leaf surface samples revealed several ASVs that were significantly associated with stomatal density (Supplemental Figure 2). We found that 40% of the resulting ASVs associated with high numbers of stomata belonged to the core microbiome that we identified here (Supplemental Table 1), whereas none of the low-stomata-associated ASVs belonged to the core microbiome.

**Figure 5:**
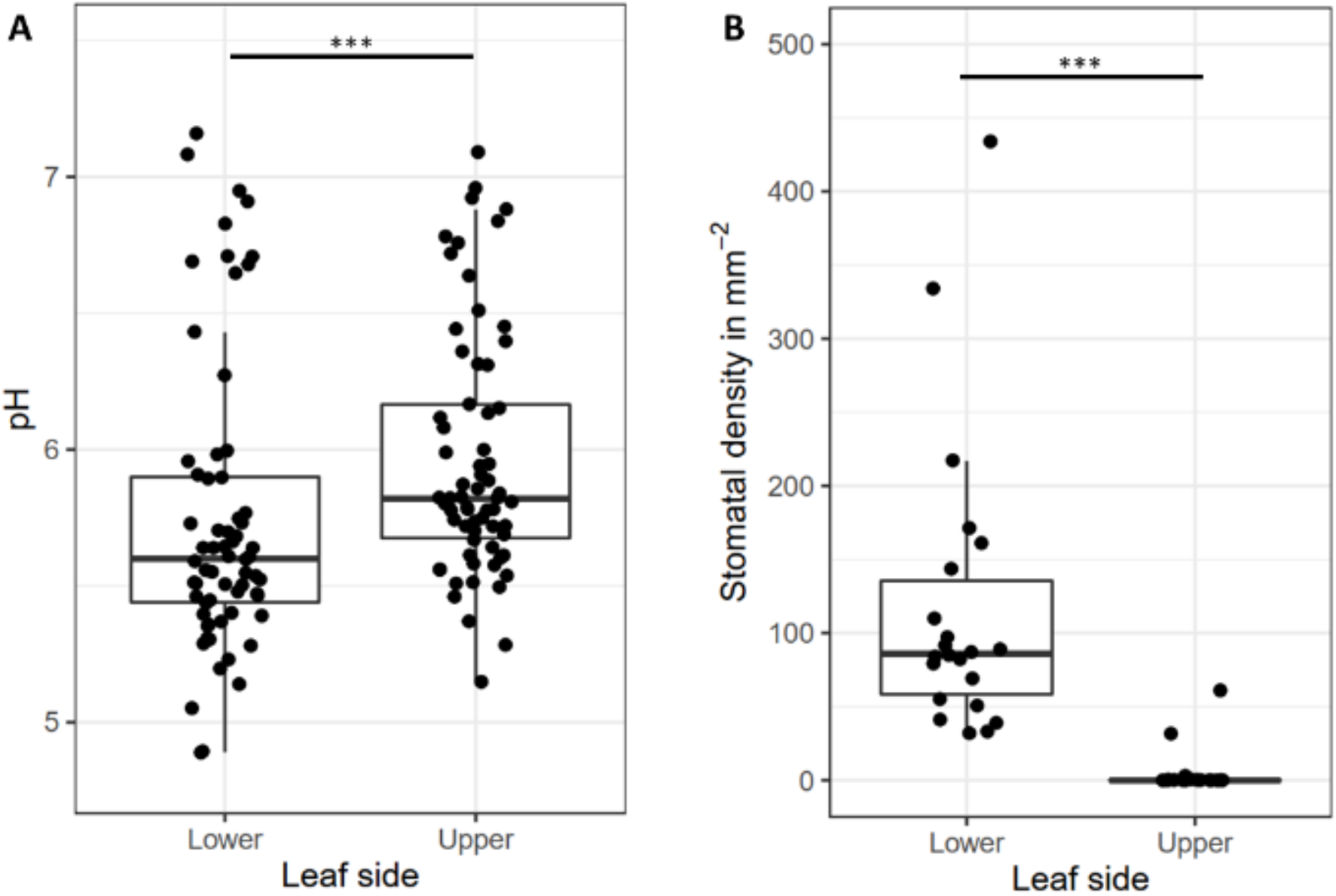
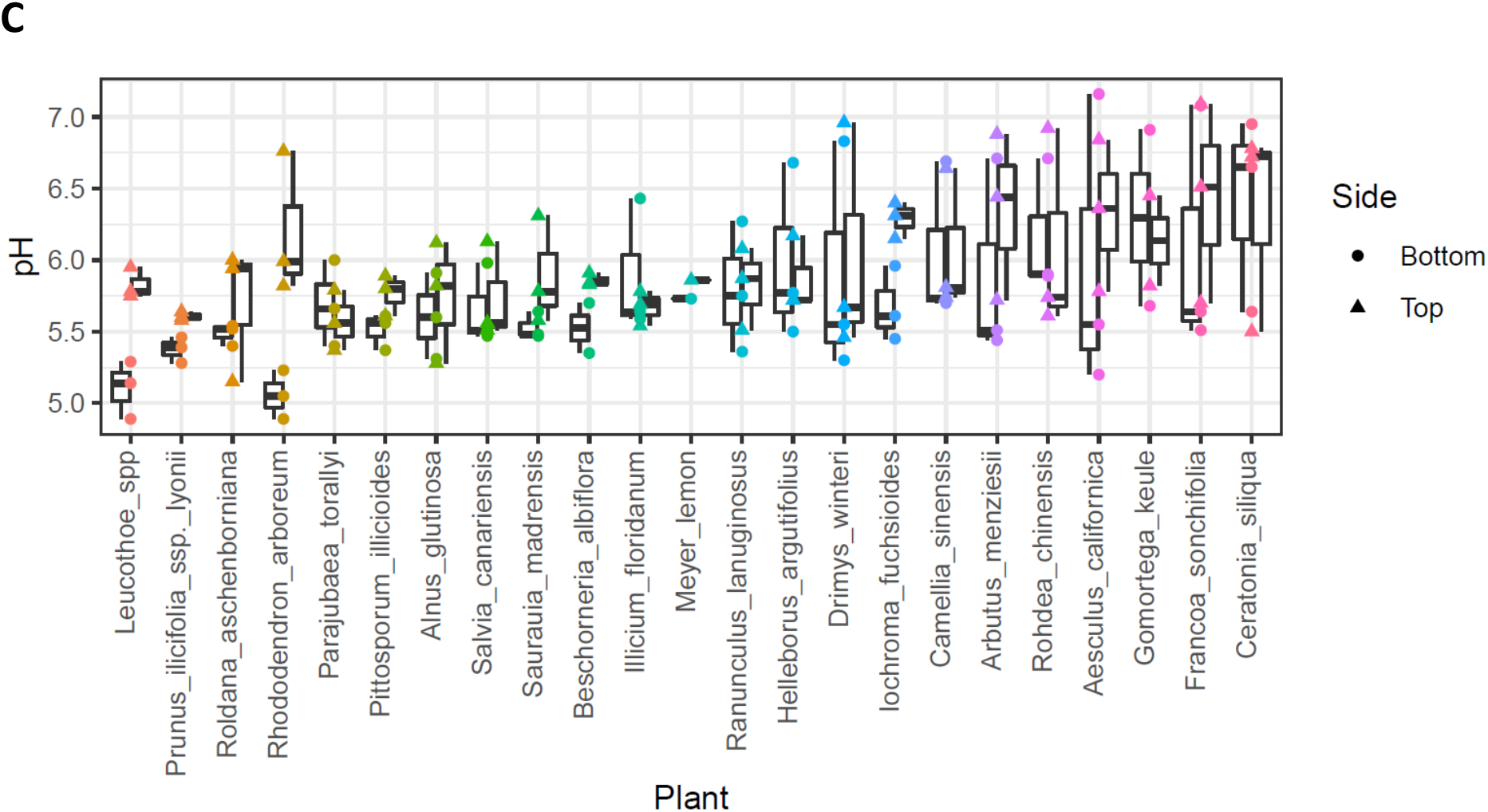
Boxplots showing mean and range of (A) pH and (B) stomatal density for lower and upper leaf sides separately (*** p-value < 0.001). (C) The pH values measured (ordinate) for each plant species (abscissa). Separate boxplots are shown for each leaf side of a given plant species.

### Lower leaf surfaces are more acidic that upper leaf surfaces

The leaf surface pH measured in this study ranged from 4.89 to 7.16. The most important leaf characteristic associated with pH was the leaf side, with the lower leaf surface being 0.21 pH units more acidic on average than the upper surface (Figure 5A). Plant species also differed significantly in pH, although the range of observed pH range of a given species was often quite high (Figure 5C). We found a significant correlation between pH and the observed ASV richness of the samples (p-value = 0.044, Kendall rank coefficient tau = 0.13), with higher - more neutral - pH values associated with higher bacterial richness, and a differential abundance analysis using DESeq2 revealed that the abundances of several bacterial ASVs were significantly correlated with pH (Figure 4A).

## Discussion

Our common garden study of the phyllosphere microbiome across 24 plant species revealed plant species identity to be the most important factor driving community composition, confirming results of previous studies (12, 20, 22, 39). When zooming in from the landscape to single-leaf level, however, we find differences in microbial community structure to be largely driven by leaf side. Upper and lower leaf surfaces differed in pH and stomatal density but were also observed to be differentially impacted by microbial dispersal. The distinction between upper and lower leaf surfaces was characterized by a strong interaction effect between plant species and leaf side (Table 1). On the lower leaf surfaces, higher host-plant-species filtering (higher R^2^ in the PERMANOVA model), lower richness (Figure 3A, B), and higher core ASV abundance (Figure 2D) indicate that this is a more selective, plant-associated environment than the upper leaf surface. On the upper leaf surfaces, ASVs were more variable (Figure 2C, E), less endemic (Figure 3C), maintained higher community richness (Figure 3A, B), and fit better to the occupancy-abundance neutral model (Figure 2A, B). These patterns suggest stochastic forces play a larger role in top side leaf community structure, due to decreased selection (plant filtering) and/or higher dispersal. Focusing on this top-bottom dichotomy is likely to yield observations that are more informative of the role of both the plant features driving phyllosphere microbial community assembly, and conversely the role of microbiomes in shaping plant functional traits (12) and ecosystem function (40).

Within this leaf side framework, our study illustrates how bacterial community structure is not only predicted by leaf side itself, but by the varying topographic (i.e. stomatal density) and chemical (i.e. surface pH) characteristics distinguished between top and bottom. Leaf surface pH was significantly correlated with leaf microbial community richness. Similar to previous results from soil (34) and wetland (35) environments, leaf surface community richness was found to increase as pH shifted towards more neutral conditions, with several taxa being significantly correlated with pH (Figure 3A). These results are perhaps unsurprising given the conservation of microbial pH preference (41) and strong effect of pH on microbial metabolism (42). However, it is notable that the impact of pH was significant in this study despite a limited observed pH range (pH 4.9 - 7.2; Figure 5A, C) compared to previous phyllosphere pH observations (pH 1 (carnivorous plants) – pH 11 (Malvaceae))(43, 44). Interestingly, top sides of leaves were significantly more pH neutral, which may allow more bacterial taxa, including non-host specific taxa, to colonize the upper leaf surfaces.

Stomata are a critical leaf trait regulating processes such as gas exchange, transpiration, and microbial defense (45). In relation to the leaf surface microbiome, studies have shown stomatal grooves aid in microbial retention during chemical disturbance (46) and to a lesser degree leaf surface dessication (47). In line with previous observations, we find stomata to exist almost exclusively on the lower leaf surfaces (Figure 5B), where stomatal density was found to be negatively correlated with microbial diversity (Figure 3A, B). Although further work is required to deliniate the cause of this pattern, we speculate that stomata act as a component of plant filtering effects in the phyllosphere, that may play act as micro-refugia or regulators of the micro-environment within the leaf boundary layer via transpiration.

Overall, despite considerable diversity in leaf morphologies and chemistries across plant species, phyllosphere microbial community structure has been traditionally overlooked at the single-leaf level in favor of macro-landscape level studies. Our study illustrates how changing the scale in which we observe microbial communities can impact our understandings of phyllosphere microbiome assembly, stability, and function.

## Materials and Methods

### Sampling location

Plants were sampled from the University of California, Berkeley Botanical Garden (Berkeley, California, U.S.) (37°52’30”N, 122°14’15”W) over four days from 14 April, 2021 to 19 April, 2021. During this period, the 6-hour temperature averages ranged from 8°C to 23°C, the relative humidity average ranged from 47% to 88%, and no precipitation occurred. However, most plants in the botanical garden were irrigated at least once a week. We selected plant species so that our samples would cover a wide range of leaf morphologies and plant growth forms (tree-like, shrub, or herbaceous). Furthermore, the leaves had to be at least two months old by the time we sampled, and each sampled plant species was preferably represented by three individuals in the botanical garden. We only sampled leaves that did not touch any soil surface to minimize soil contamination. Due to limits of swab sampling, conifers and plants with very small leaves were not included. In total, we sampled 24 different plant species each of which three individuals were sampled randomly over the different days (due to practical reasons four species were represented by only one or two individuals).

### Leaf sampling

Leaves were excised by cutting off the petiole with shears sterilized with 70% ethanol, immediately placed in a sterile plastic bag, and processed within one hour of sampling. To provide a sufficient area for microbial and pH sampling, 1 to 8 leaves, depending on leaf sizes, corresponding to a total leaf area ranging from 94 to 276 cm^2^, were sampled from each individual plant. The coordinates of each sampled plant were obtained using a Garmin eTrex 30x GPS device.

### Microbial measurements

To sample the microbiome on leaf surfaces, a sterile flock swab (Puritan, 25-3306-H) was dipped in a sterile wash solution (autoclaved 10 mM MgCl_2_ containing 0.002% Tween). It was then used to thoroughly swab one half of a leaf surface (left of midrib, including half of the midrib) of all leaves. Swabs were then placed in sterile, dry 1.5 mL tubes, transported to the lab on ice and stored at −20°C the same day. One blank swab, dipped in wash solution and placed directly in a 1.5 mL tube was included as a control. For each plant sample, the lower and upper sides of the leaf were swabbed separately. For each sampled leaf, the left half of both the lower and upper sides were separately swabbed, allowing pH measurements to be taken of the non-disturbed, right half of each leaf.

### pH measurements

Following swabbing to remove bacteria from the leaf surface, the pH of the upper and lower leaf surface was measured using a sterile cotton swab (Puritan, 806-WC). Briefly, a swab was soaked in a 1.5 mL tube containing 300μL of sterile deionized H_2_O, the right halves of the leaves were swabbed with the wetted swab and the swab was then returned to the 1.5 mL tube and the wooden handle excised to ensure the tube could be closed. The tube was vortexed with the swab for 20s. The swab was then pressed against the sides of the tube to squeeze liquid out and discarded. The pH probe (Sartorius, PY-P21) was placed in the small volume in the tube, and pH meter (Denver instruments, UB-5) was used to measure pH after readings stabilized within 2 minutes. Consistency of the leaf surface pH methodology was based on preliminary trials across various plant species. These trials revealed that our chosen methodology yielded pH values that were similar to that obtained by placing a flat-bottom electrode on a wetted leaf for 15 minutes, a method used for leaf pH measurements previouslyJ. Oertli et al. (48).

### Leaf characteristics

Following measurement of pH, several leaf characteristics, including stomatal density, leaf hardness, and leaf pubescence, were recorded. Stomatal density was measured using epidermal leaf impressions (49). Temporary stomatal impressions were made by coating unswabbed leaf samples with a layer of clear nail varnish. For each leaf, stomatal slides were taken near the midvein at the base, middle, and tip of each leaf, avoiding the midrib. After drying, impressions were removed, observed, and imaged under a light microscope at 200 X (Olympus, BH-2; Nikon, D200). All stomata in one microscopic field were counted, and stomatal density was determined as the mean of the three leaf locations, after normalizing for the magnification and area examined. Leaf area was measured using an area meter (LI-COR, LI-3100C). Leaf hardness was judged on a qualitative scale of zero to three, with zero being soft/thin and four being hard/brittle (50). Lastly, leaf pubescence was assessed based on the presence or absence of leaf hairs visualized under a microscope.

### DNA extraction and sequencing

Microbial swabs were sent to the University of Colorado, Boulder for DNA extraction and marker gene sequencing. DNA was extracted from the swabs using a tube-based DNeasy Powersoil Pro Kit (Qiagen) to minimize the occurrence of well-to-well contamination of the presumably low-biomass leaf surface samples. DNA extractions were performed according to manufacturer protocol with two exceptions: after CD1 addition and before the bead beating step, samples were incubated at 65°C for 10 minutes; and DNA was eluted with 50 μL C6 solution as opposed to 100 μL.

All leaf swab DNA extractions were PCR amplified targeting the 16S rRNA gene (hypervariable V4–V5 region) using 515f/806r barcoded primers (51, 52). PCR amplification was performed in duplicate for 133 leaf swab samples, 14 DNA extraction blank negative controls, and 13 no-template control (PCR) negative controls. Resulting amplicons were pooled, cleaned, and normalized using SequalPrep Normalization plates (Thermo Fisher Scientific, Waltham, MA). Pooled samples were sent to CU Anschutz School of Medicine core facility for Next Generation Sequencing on an Illumina MiSeq using 2×150 paired end chemistry.

### Bioinformatic analyses

Amplicon Sequence Variants (ASVs) were generated from raw reads using an in-house bioinformatic pipeline based around DADA2 (53) as performed previously (DADA2 version 1.14.1, Fierer Lab DADA2 Pipeline Version 0.1.0) (54). Prior to sequence inference, reads were filtered and trimmed using the following settings: truncLen=c(145,150), maxEE=c(2,2), truncQ=2, maxN=0, rm.phix=TRUE. Reads from all samples were pooled for sequence inference using the pool = TRUE parameter of the dada() function. The resulting ASVs were further processed to remove chimeras and assign taxonomy using the DADA2 naïve Bayesian classifier with the SILVA database v 132 (55).

Data were imported in R and further processing was done in RStudio using R version 3.6.3 mainly using the R packages tidyverse (56) and tidyamplicons (https://github.com/Swittouck/tidyamplicons). Reads classified as chloroplasts, mitochondrial and non-bacterial were removed from the dataset. From blanks and no-template controls, we identified 15 ASVs as contaminants using the R package decontam (57) and discarded these contaminants from the dataset (losing 0.05% of the reads). In addition, in each sample we removed ASVs represented by two or fewer reads. Only when stated for specific analyses, data were normalized by rarefying samples to 4000 reads or analyzing relative abundances, in all other analyses, read counts were used.

### Statistical analyses

Statistical analyses were done using R, mainly using R package vegan (58). Results were considered significant at a p-value < 0.05. The core ASVs were determined using the method based on abundance-occupancy distributions as described by A. Shade and N. Stopnisek (38) on the rarefied dataset (Supplemental code). Alpha diversity, observed ASV richness and the inverse Simpson index, were also determined using the rarefied dataset. Differences between means were tested using the wilcox.test function as data were not normally distributed, and correlations between factors were determined using the Kendall rank correlation test.

Beta dissimilarities between samples were determined as Bray-Curtis dissimilarities based on ASV relative abundances within samples. We used PERMANOVA models to identify factors explaining variation in the Bray-Curtis dissimilarities. In these models, sample characteristics were added in order of specificity, with the most specific characteristics (e.g. stomatal density) first and the factors encompassing more characteristics (e.g. plant species) later, to avoid that the latter partly includes the effect of the more specific characteristics. The order of more general factors like region of origin and plant sample were ordered according to their nestedness, allowing PERMANOVA to determine an R^2^ value for each of these factors.

To determine which taxa were correlated with pH, stomatal density, and leaf side, we used differential abundance analyses of the R package DESeq2 (59). Due to the requirement of the original read count data in this analysis, we started from the unrarefied dataset where ASVs represented by one or two reads per sample were not discarded but ASVs that had a total relative abundance of less than 0.1% were discarded. For the continuous variables, stomatal density and pH, we did an analysis using the likelihood ratio test (LRT) of the DESeq function. For leaf side, we did a Wald significance test of the DESeq function.

The degree of endemism of each ASV was determined, after discarding ASVs that occurred in no more than one sample. Here, as a proxy for degree of endemism of an ASV, we determined the proportion of plant species on which it was not detected. Weighted endemism of samples was then calculated as the product of the relative abundance of an ASV and its degree of endemism. We log-transformed weighted endemism to analyze normally distributed data. We then optimized a multiple regression model including pH, leaf side, and stomatal density as explanatory variables for weighted endemism of samples. We used a paired Wilcoxon test to assess differences of endemism between upper and lower surfaces of leaves.

To assess correlation of host phylogeny and microbial community structure, we determined phylogenetic distances for all plant species sampled in this study and for all bacterial ASVs recovered in this study. For the ASVs, we built an approximate maximum likelihood tree based on a generalized time-reversible model. First, sequences of all ASVs were aligned using MUSCLE (60). We then used FastTree (version 2.1.9, (61)) with the Generalized Time-Reversible and nucleotide alignment specifications (-gtr -nt) to generate a phylogenetic tree. The resulting phylogenetic distances were used to determine weighted unifrac distances among samples (62). Additionally, we constructed a phylogenetic tree for all plant species used in this study. Sequences of plant species were obtained from publicly available datasets in NCBI genbank. Multiple sequence alignment of 18S rRNA genes was performed using Unipro UGENE (63). The aligned sequences were manually adjusted when necessary. Gaps and ambiguously aligned nucleotide positions were excluded from the datasets. A phylogenetic tree for plant species was constructed using maximum-likelihood method in R version 4.0.12 under the GTR+I model of evolution with 100 bootstrap replicates, using packages ape version 5.4-1 (64) and ggtree version 1.0 (65). After determining phylogenetic distances between plant hosts and between ASVs, we did a host-parasite association test as described by P. Legendre et al. (66) for the 1000 most abundant ASVs in our dataset.

## Acknowledgements

We would like to acknowledge the contribution of the UC Berkeley Botanical Garden staff and volunteers, especially Holly Forbes, Clare Loughran, and Gideon Dollarhide whose expertise was essential in choosing and identifying plant species to sample. We thank Fernando Diaz, an NSF REPS fellow (NSF grant # 1942881), for their aid in sampling. We furthermore acknowledge funding supporting this research, including a BAEF fellowship and FWO grant to WS, an NSF REPS fellowship (NSF grant # 1942881) to FTD, a Ford fellowship to MKC, an NSF GRF to CMW, and funding from the U.S. National Science Foundation (G-03583-01) to NF. BK is a Chan Zuckerberg Biohub Investigator and a fellow at the Wissenschaftskolleg zu Berlin.

## Supplemental material

**Supplemental Figure 1:**
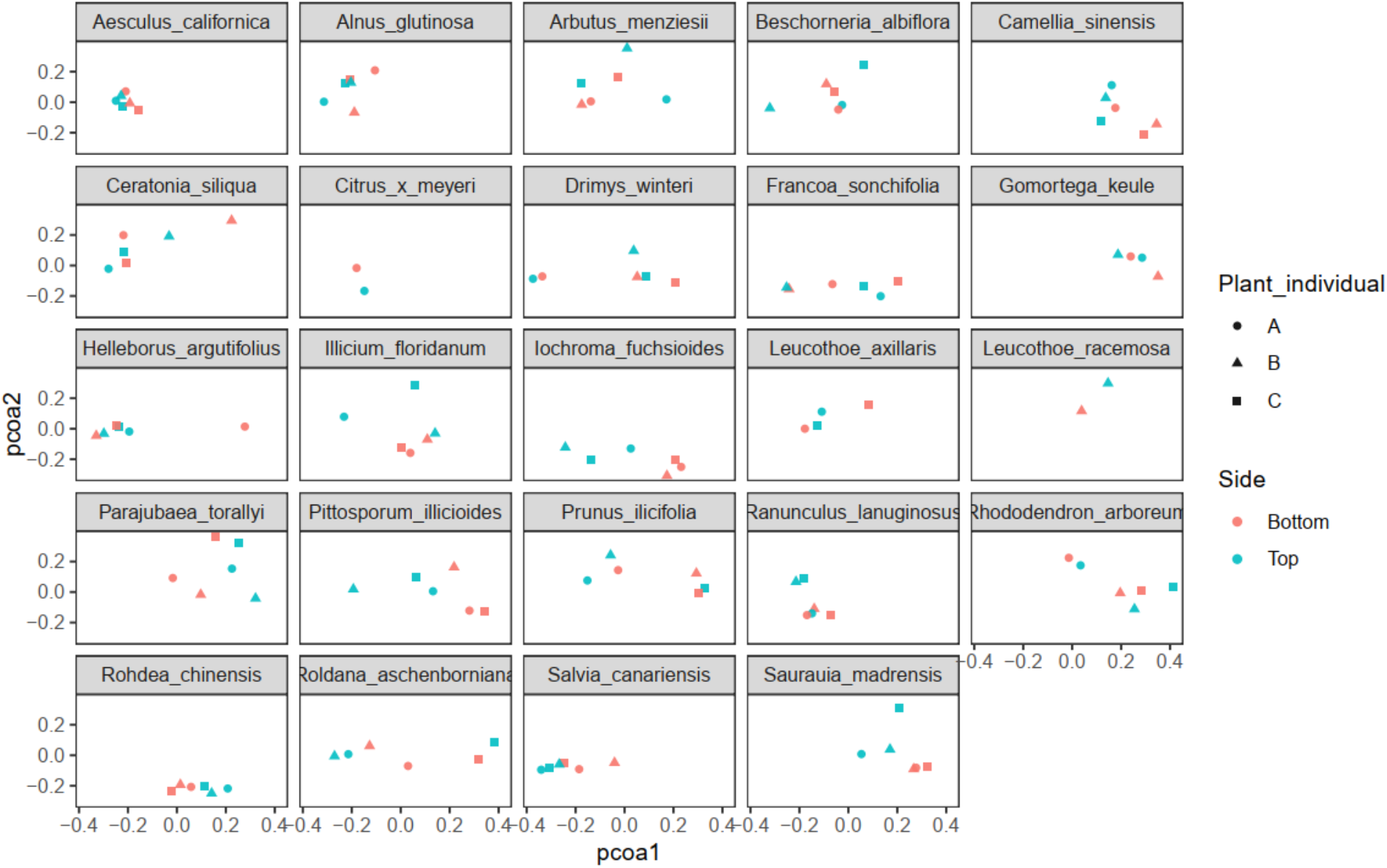
Principal coordinate analysis (PCoA) of all samples, with samples represented per plant species in each panel.

**Supplemental Figure 2:**
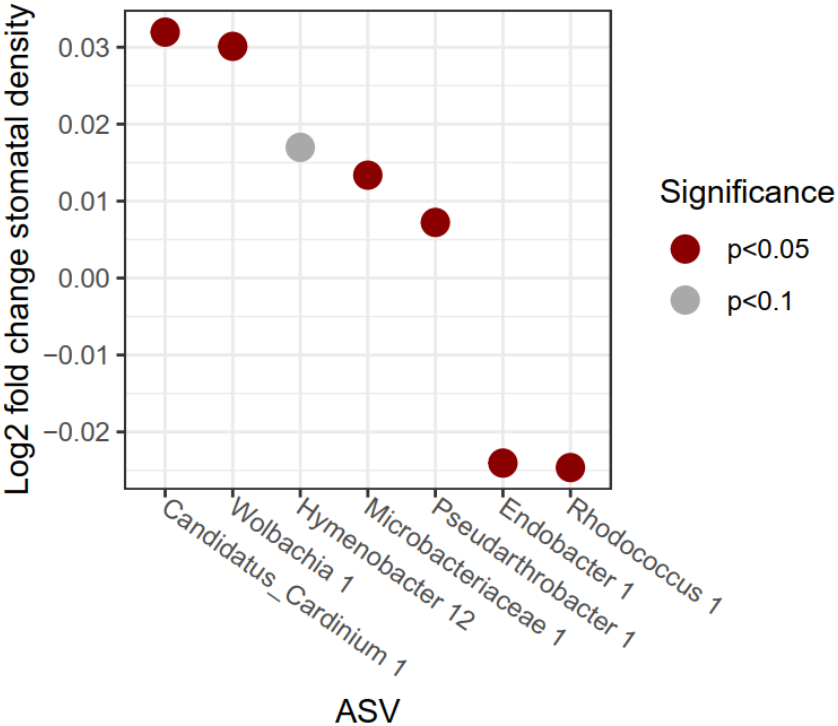
ASVs that linearly correlated with the log of stomatal density (mm^-2^) on the lower leaf surface.

**Supplemental Table 1:**
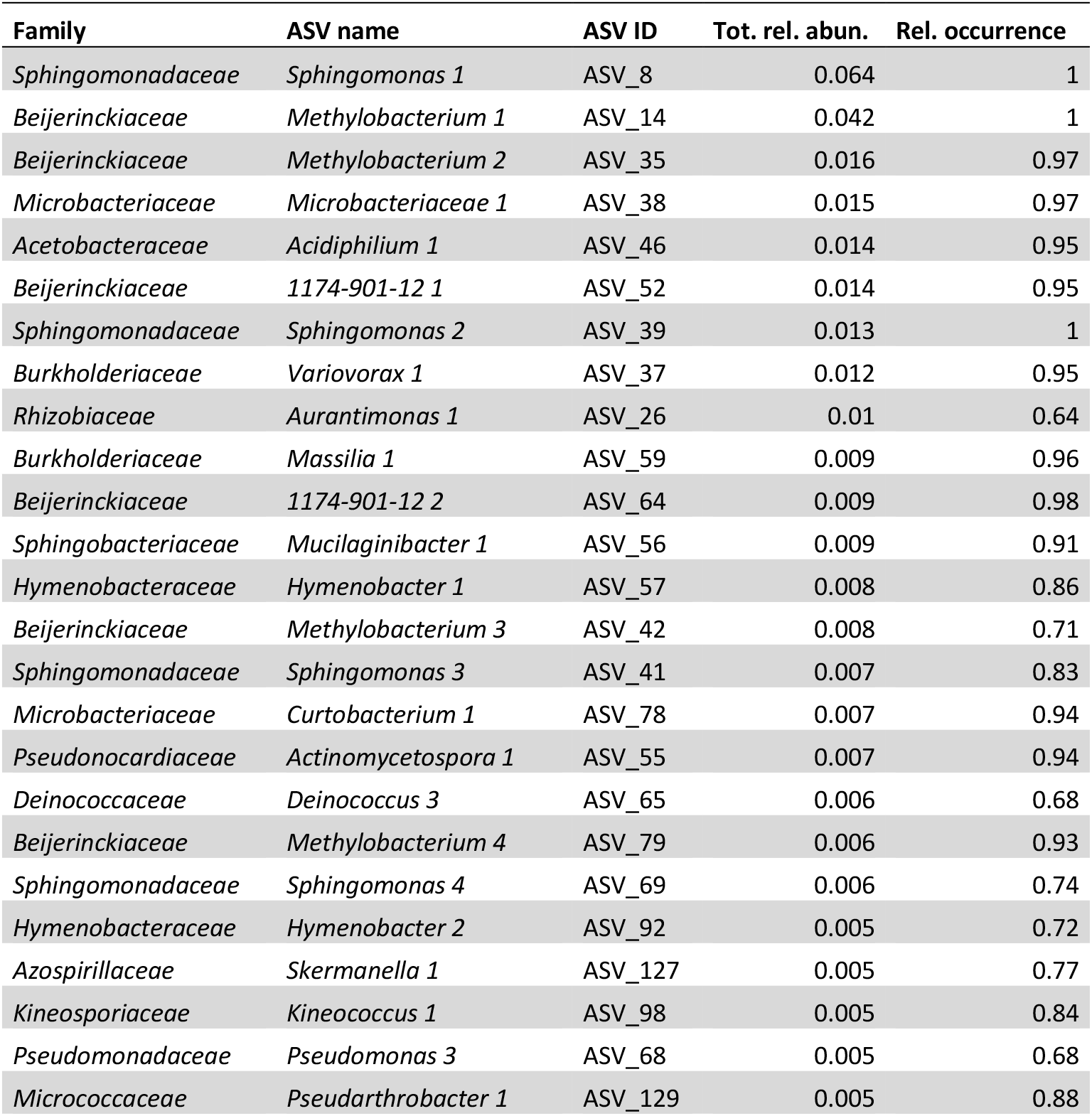
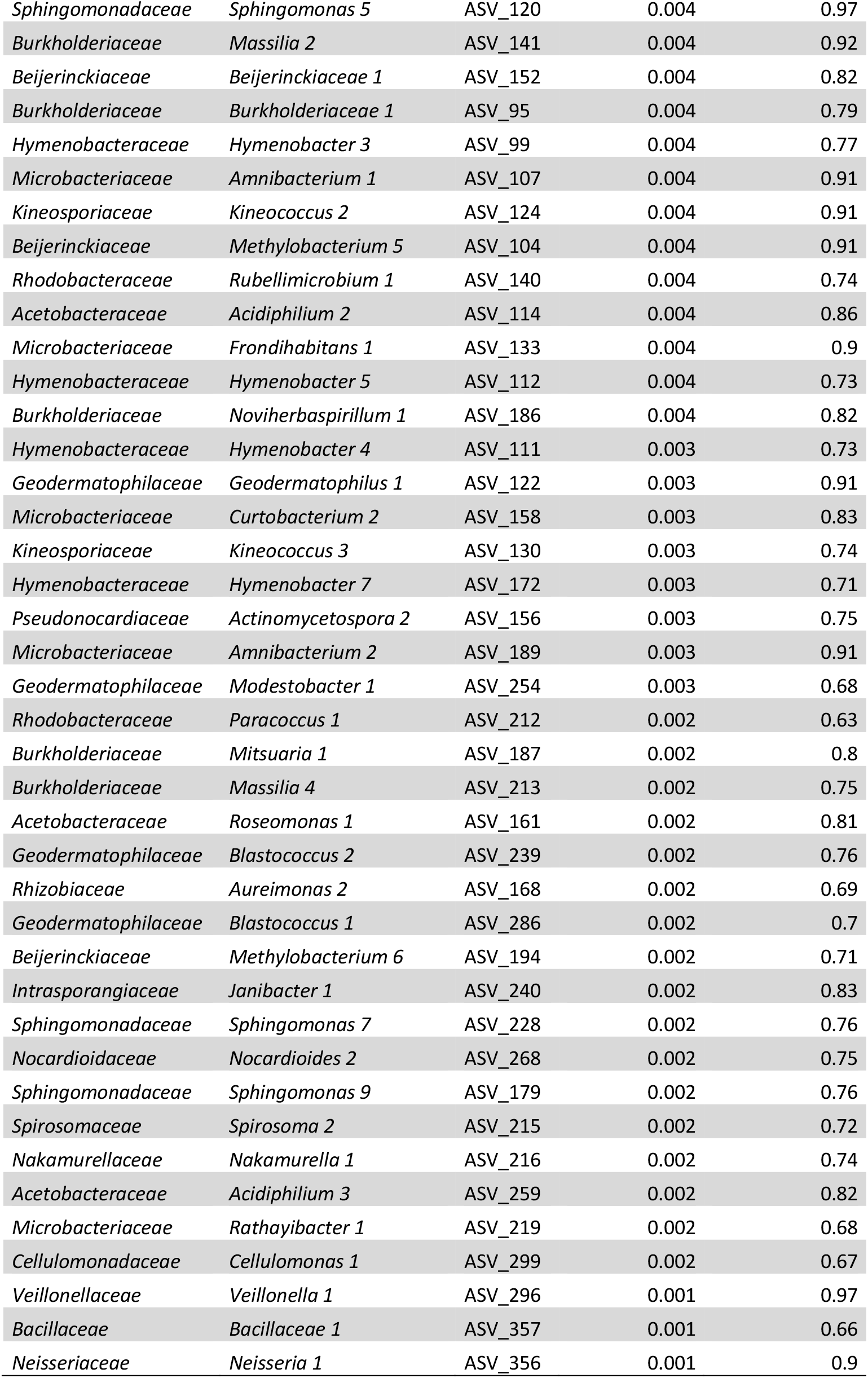
Core phyllosphere microbiome on plants collected at the UC Berkeley botanical garden (California, USA). Core ASVs were determined based on abundance-occupancy distributions and their contribution to beta diversity (38). The total relative abundance of an ASV in the dataset and the proportion of samples in which it was found are included.

| Family | ASV name | ASV ID | Tot. rel. abund. | Rel. occurrence |
| --- | --- | --- | --- | --- |
| *Sphingomonadaceae* | *Sphingomonas* 1 | ASV_8 | 0.064 | 1 |
| *Beijerinckiaceae* | *Methylobacterium* 1 | ASV_14 | 0.042 | 1 |
| *Beijerinckiaceae* | *Methylobacterium* 2 | ASV_35 | 0.016 | 0.97 |
| *Microbacteriaceae* | *Microbacteriaceae* 1 | ASV_38 | 0.015 | 0.97 |
| *Acetobacteraceae* | *Acidiphilium* 1 | ASV_46 | 0.014 | 0.95 |
| *Beijerinckiaceae* | 1174-901-12 1 | ASV_52 | 0.014 | 0.95 |
| *Sphingomonadaceae* | *Sphingomonas* 2 | ASV_39 | 0.013 | 1 |
| *Burkholderiaceae* | *Variovorax* 1 | ASV_37 | 0.012 | 0.95 |
| *Rhizobiaceae* | *Aurantimonas* 1 | ASV_26 | 0.01 | 0.64 |
| *Burkholderiaceae* | *Massilia* 1 | ASV_59 | 0.009 | 0.96 |
| *Beijerinckiaceae* | 1174-901-12 2 | ASV_64 | 0.009 | 0.98 |
| *Sphingobacteriaceae* | *Mucilaginibacter* 1 | ASV_56 | 0.009 | 0.91 |
| *Hymenobacteraceae* | *Hymenobacter* 1 | ASV_57 | 0.008 | 0.86 |
| *Beijerinckiaceae* | *Methylobacterium* 3 | ASV_42 | 0.008 | 0.71 |
| *Sphingomonadaceae* | *Sphingomonas* 3 | ASV_41 | 0.007 | 0.83 |
| *Microbacteriaceae* | *Curtobacterium* 1 | ASV_78 | 0.007 | 0.94 |
| *Pseudonocardiaceae* | *Actinomycetospora* 1 | ASV_55 | 0.007 | 0.94 |
| *Deinococcaceae* | *Deinococcus* 3 | ASV_65 | 0.006 | 0.68 |
| *Beijerinckiaceae* | *Methylobacterium* 4 | ASV_79 | 0.006 | 0.93 |
| *Sphingomonadaceae* | *Sphingomonas* 4 | ASV_69 | 0.006 | 0.74 |
| *Hymenobacteraceae* | *Hymenobacter* 2 | ASV_92 | 0.005 | 0.72 |
| *Azospirillaceae* | *Skermanella* 1 | ASV_127 | 0.005 | 0.77 |
| *Kineosporiaceae* | *Kineococcus* 1 | ASV_98 | 0.005 | 0.84 |
| *Pseudomonadaceae* | *Pseudomonas* 3 | ASV_68 | 0.005 | 0.68 |
| *Micrococcaceae* | *Pseudarthrobacter* 1 | ASV_129 | 0.005 | 0.88 |
| *Sphingomonadaceae* | *Sphingomonas* 5 | ASV_120 | 0.004 | 0.97 |
| *Burkholderiaceae* | *Massilia* 2 | ASV_141 | 0.004 | 0.92 |
| *Beijerinckiaceae* | *Beijerinckiaceae* 1 | ASV_152 | 0.004 | 0.82 |
| *Burkholderiaceae* | *Burkholderiaceae* 1 | ASV_95 | 0.004 | 0.79 |
| *Hymenobacteraceae* | *Hymenobacter* 3 | ASV_99 | 0.004 | 0.77 |
| *Microbacteriaceae* | *Amnibacterium* 1 | ASV_107 | 0.004 | 0.91 |
| *Kineosporiaceae* | *Kineococcus* 2 | ASV_124 | 0.004 | 0.91 |
| *Beijerinckiaceae* | *Methylobacterium* 5 | ASV_104 | 0.004 | 0.91 |
| *Rhodobacteraceae* | *Rubellimicrobium* 1 | ASV_140 | 0.004 | 0.74 |
| *Acetobacteraceae* | *Acidiphilium* 2 | ASV_114 | 0.004 | 0.86 |
| *Microbacteriaceae* | *Fronidhabitans* 1 | ASV_133 | 0.004 | 0.9 |
| *Hymenobacteraceae* | *Hymenobacter* 5 | ASV_112 | 0.004 | 0.73 |
| *Burkholderiaceae* | *Noviherbaspirillum* 1 | ASV_186 | 0.004 | 0.82 |
| *Hymenobacteraceae* | *Hymenobacter* 4 | ASV_111 | 0.003 | 0.73 |
| *Geodermatophilaceae* | *Geodermatophilus* 1 | ASV_122 | 0.003 | 0.91 |
| *Microbacteriaceae* | *Curtobacterium* 2 | ASV_158 | 0.003 | 0.83 |
| *Kineosporiaceae* | *Kineococcus* 3 | ASV_130 | 0.003 | 0.74 |
| *Hymenobacteraceae* | *Hymenobacter* 7 | ASV_172 | 0.003 | 0.71 |
| *Pseudonocardiaceae* | *Actinomycetospira* 2 | ASV_156 | 0.003 | 0.75 |
| *Microbacteriaceae* | *Amnibacterium* 2 | ASV_189 | 0.003 | 0.91 |
| *Geodermatophilaceae* | *Modestobacter* 1 | ASV_254 | 0.003 | 0.68 |
| *Rhodobacteraceae* | *Paracoccus* 1 | ASV_212 | 0.002 | 0.63 |
| *Burkholderiaceae* | *Mitsuaria* 1 | ASV_187 | 0.002 | 0.8 |
| *Burkholderiaceae* | *Massilia* 4 | ASV_213 | 0.002 | 0.75 |
| *Acetobacteraceae* | *Roseomonas* 1 | ASV_161 | 0.002 | 0.81 |
| *Geodermatophilaceae* | *Blastococcus* 2 | ASV_239 | 0.002 | 0.76 |
| *Rhizobiaceae* | *Aureimonas* 2 | ASV_168 | 0.002 | 0.69 |
| *Geodermatophilaceae* | *Blastococcus* 1 | ASV_286 | 0.002 | 0.7 |
| *Beijerinckiaceae* | *Methylobacterium* 6 | ASV_194 | 0.002 | 0.71 |
| *Intrasporangiaceae* | *Janibacter* 1 | ASV_240 | 0.002 | 0.83 |
| *Sphingomonadaceae* | *Sphingomonas* 7 | ASV_228 | 0.002 | 0.76 |
| *Nocardioideae* | *Nocardioideae* 2 | ASV_268 | 0.002 | 0.75 |
| *Sphingomonadaceae* | *Sphingomonas* 9 | ASV_179 | 0.002 | 0.76 |
| *Spirosomaceae* | *Spirosoma* 2 | ASV_215 | 0.002 | 0.72 |
| *Nakamurellaceae* | *Nakamurella* 1 | ASV_216 | 0.002 | 0.74 |
| *Acetobacteraceae* | *Acidiphilium* 3 | ASV_259 | 0.002 | 0.82 |
| *Microbacteriaceae* | *Rathayibacter* 1 | ASV_219 | 0.002 | 0.68 |
| *Cellulomonadaceae* | *Cellulomonas* 1 | ASV_299 | 0.002 | 0.67 |
| *Veillonellaceae* | *Veillonella* 1 | ASV_296 | 0.001 | 0.97 |
| *Bacillaceae* | *Bacillaceae* 1 | ASV_357 | 0.001 | 0.66 |
| *Neisseriaceae* | *Neisseria* 1 | ASV_356 | 0.001 | 0.9 |

**Supplemental Table 2:** PERMANOVA of all factors included in this study.

| Factor | R<sup>2</sup> | p-value |
| --- | --- | --- |
| Leaf surface pH | 0.02 | <0.001 |
| Stomatal density | 0.02 | <0.001 |
| Leaf side (upper/lower) | 0.02 | <0.001 |
| Leaf pubescence | 0.04 | <0.001 |
| Leaf hardness | 0.03 | <0.001 |
| Evergreen or deciduous plant | 0.03 | <0.001 |
| Plant type (herb/shrub/tree) | 0.04 | <0.001 |
| Climate of native origin | 0.06 | <0.001 |
| Region of native origin | 0.11 | <0.001 |
| Host plant species | 0.10 | <0.001 |
| Date sampled | 0.03 | <0.001 |
| Plant individual / leaf sample | 0.21 | <0.001 |
| Interaction of pH and host plant species | 0.17 | 0.001 |
| Interaction of side and host plant species | 0.11 | 0.016 |

